# The mitochondrial thiolase ACAT1 regulates monocyte/macrophage type I interferon *via* epigenetic control

**DOI:** 10.1101/2024.01.29.577773

**Authors:** Jing Wu, Komudi Singh, Vivian Shing, Anand K. Gupta, Rebecca D. Huffstutler, Duck-Yeon Lee, Michael N. Sack

## Abstract

Lipid-derived acetyl-CoA is shown to be the major carbon source for histone acetylation. However, there is no direct evidence demonstrating lipid metabolic pathway contribututions to this process. Mitochondrial acetyl-CoA acetyltransferase 1 (ACAT1) catalyzes the final step of ß-oxidation, the aerobic process catabolizing fatty acids (FA) into acetyl-CoA. To investigate this in the context of immunometabolism, we generated macrophage cell line lacking ACAT1. ^13^C-carbon tracing combined with mass spectrometry confirmed incorporation of FA-derived carbons into histone H3 and this incorporation was reduced in ACAT1 KO macrophage cells. RNA-seq identified a subset of genes downregulated in ACAT1 KO cells including STAT1/2 and interferon stimulated genes (ISGs). CHIP analysis demonstrated reduced acetyl-H3 binding to STAT1 promoter/enhancer regions. Increasing histone acetylation rescued STAT1/2 expression in ACAT1 KO cells. Concomitantly, ligand triggered IFNβ release was blunted in ACAT1 KO cells and rescued by reconstitution of ACAT1. Furthermore, ACAT1 promotes FA-mediated histone acetylation in an acetylcarnitine shuttle-dependent manner. In patients with obesity, levels of ACAT1 and histone acetylation are abnormally elevated. Thus, our study identified a novel link between ACAT1 mediated FA metabolism and epigenetic modification on STAT1/2 that uncovers a regulatory role of lipid metabolism in innate immune signaling and opens novel avenues for interventions in human diseases such as obesity.

## INTRODUCTION

Immune cells have varying metabolic demands, based on their proliferative capacity, polarization, and activation status. In this context, it is now well established that cellular metabolism is highly integrated with immune cell fate and function. The interaction between metabolism and immune function is defined as immunometabolism and encompasses: (i) energy substrate utilization (fate-dependent metabolic remodeling) (1–3); (ii) the role of metabolic intermediates as signaling molecules (4–6); and (iii) by the indirect sequelae of metabolism on intracellular organelle function and fidelity resulting in subsequent retrograde signaling (7–9).

Metabolic remodeling in conjunction with altered immune cell fates is most well characterized and has been extensively reviewed (1–3). The metabolites most well characterized that function as signaling intermediates governing immune cell fate include tricarboxylic acid cycle (TCA) intermediates (4, 5, 9, 10), ketone bodies and short-chain fatty acids (SCFA’s) (11). In myeloid cells, the accumulation of TCA intermediates confers immunoregulatory effects *via* signaling through nitric oxide, reactive oxygen species and prostaglandin-mediated signaling, *via* G-protein coupled receptors, *via* activation of transcriptional regulators, *via* altering mitochondrial function and fidelity (4, 9) and by indirect effects through posttranslational modifications (6). Ketone bodies confer control on innate immune function by disrupting inflammasome complex formation (12) and or *via* the metabolic effects that modulate immune reactivity (13). Emerging evidence implicates pentose phosphate pathway and nucleotide synthesis intermediates (14), amino acids, long-chain saturated fatty acids and amino- and fatty-acid conjugates (15) in conferring immunoregulation. Finally, nutrient sensing organelles implicated in immune retrograde signaling include mitochondria (8, 16, 17), autophagosomes (14, 18, 19) and the endosome-lysosome system (20, 21).

Further investigation into the intersection between metabolic pathways, metabolic intermediates and intracellular organelle initiated signaling should provide additional insight into how modifying metabolism may influence immune cell function. To investigate the composite of these various components of immunometabolism, we focused on exploring the potential immunoregulatory role of the mitochondrial thiolase acetyl-CoA acetyltransferase 1 (ACAT1). As its name implicates, ACAT1 is mitochondrial enriched, functions as a mitochondrial acetyltransferase enzyme (22), and is a distal enzyme in the catabolism of isoleucine (23), fatty acids (24) and ketone bodies (25) to generate acyl-CoA moieties and CoA. Moreover, lipids have been shown to be a major carbon source for histone acetylation (26), however the contribution of fatty acid oxidation (FAO) enzymes to this process has not been explored and reported. We postulated, that the study of this enzyme may shed additional insight into how mitochondrial metabolism and signaling controls myeloid immune cell function.

In this study, we find that type I IFN interferon (T1IFN) signaling was blunted in myeloid cells following the genetic depletion of ACAT1. Additionally, this effect was due to the reduction in fatty acid orchestrated histone acetylation and attenuation in transcript and protein levels of a canonical T1IFN regulatory protein STAT1. This fatty acid catabolism effect was confirmed in that the modulation of histone acetylation is dependent on the carnitine/acylcarnitine translocase. Finally, in human subjects with obesity, circulating monocytes express excessive ACAT1 compared to lean control subjects, and this is associated with increased histone acetylation and increased transcript levels of canonical T1IFN pathway genes. Taken together, these data support that ACAT1-mediated fatty acid oxidation, *via* increased histone acetylation, orchestrate the induction of myeloid type 1 interferon signaling.

## RESULTS

### ACAT1 Depletion Dampens Type I Interferon Signaling

Prior to exploring the role of ACAT1, we turned to public databases to evaluate its expression profiling in circulating immune cell. The human PBMC single cell RNA-seq data was interrogated in the immunological genome project database (https://www.immgen.org/). Analysis of the Leiden algorithm shows that Acat1 expression is highly enriched across all lineages in peripheral blood mononuclear cells (PMBCs) (Suppl. Figure 1A). Moreover, the human protein atlas (https://www.proteinatlas.org/) shows that ACAT1 is localized to mitochondria and expressed across PBMC cell populations with lower levels in polymorphonuclear leukocytes (PMNs) (Suppl. Figure 1B). To explore the role of ACAT1 in innate immune cells, we employed the murine J774A.1 macrophage cell line and generated isogenic cell line lacking ACAT1 *via* CRISPR-Cas9 gene editing. RNA-seq analysis was then performed on the unstimulated or LPS-stimulated WT *vs* ACAT1 KO macrophages. PCA plots of the differentially expressed (DE) genes showed that the presence/absence of Acat1 gene was the principle component driving gene expression changes, irrespective of the cell activation status (Suppl. Figure 2). 3288 genes were differentially expressed with 1618 genes being induced and 1670 being reduced in unstimulated ACAT1 KO cells (Suppl. Table 1). Cluster Profiler pathway enrichment analysis curated for pathways linked to immune cell type and signaling showed that the most highly regulated DE genes were linked to the response to viral infections and in response to interferon beta (IFN-β) (Figure 1A). As depicted on the volcano plot of DE genes, transcripts encoding signaling and regulatory proteins in Type I IFN (T1IFN) were robustly downregulated following Acat1 deletion (Figure 1B). Quantitative RT-PCR confirmed the reduction in transcript levels of T1IFN pathway genes in ACAT1^-/-^ (KO) macrophages (Figure 1C). To functionally validate these transcript signatures, WT and ACAT1 KO cells were stimulated by various T1IFN inducers, including lipopolysaccharide (LPS) (TLR4 agonist), 2’,3’-cGAMP (STING agonist) and poly(I:C) (TLR3 agonist). Here, the levels of IFN-β secretion were diminished following ablation of Acat1 compared to WT control (Figure 1, D and E). Interestingly, the canonical TLR4 induced cytokine, TNFα was not altered comparing WT *vs* ACAT1 KO cells (Figure 1D, right panel). We then assessed canonical T1IFN signaling in these cells in response to LPS and found that activation of this pathway including phosphorylation of STAT1 and STAT2 were markedly blunted in Acat1 KO cells. Notably, the steady state levels of STAT1 and STAT2 were significantly reduced, while interferon regulatory factor 3 (IRF3) levels were unchanged (Figure 1F). Together, these data implicated a vital role of ACAT1 in the regulation of type I IFN response primarily by the alteration of regulatory control of selective gene expression.

**Figure 1.**
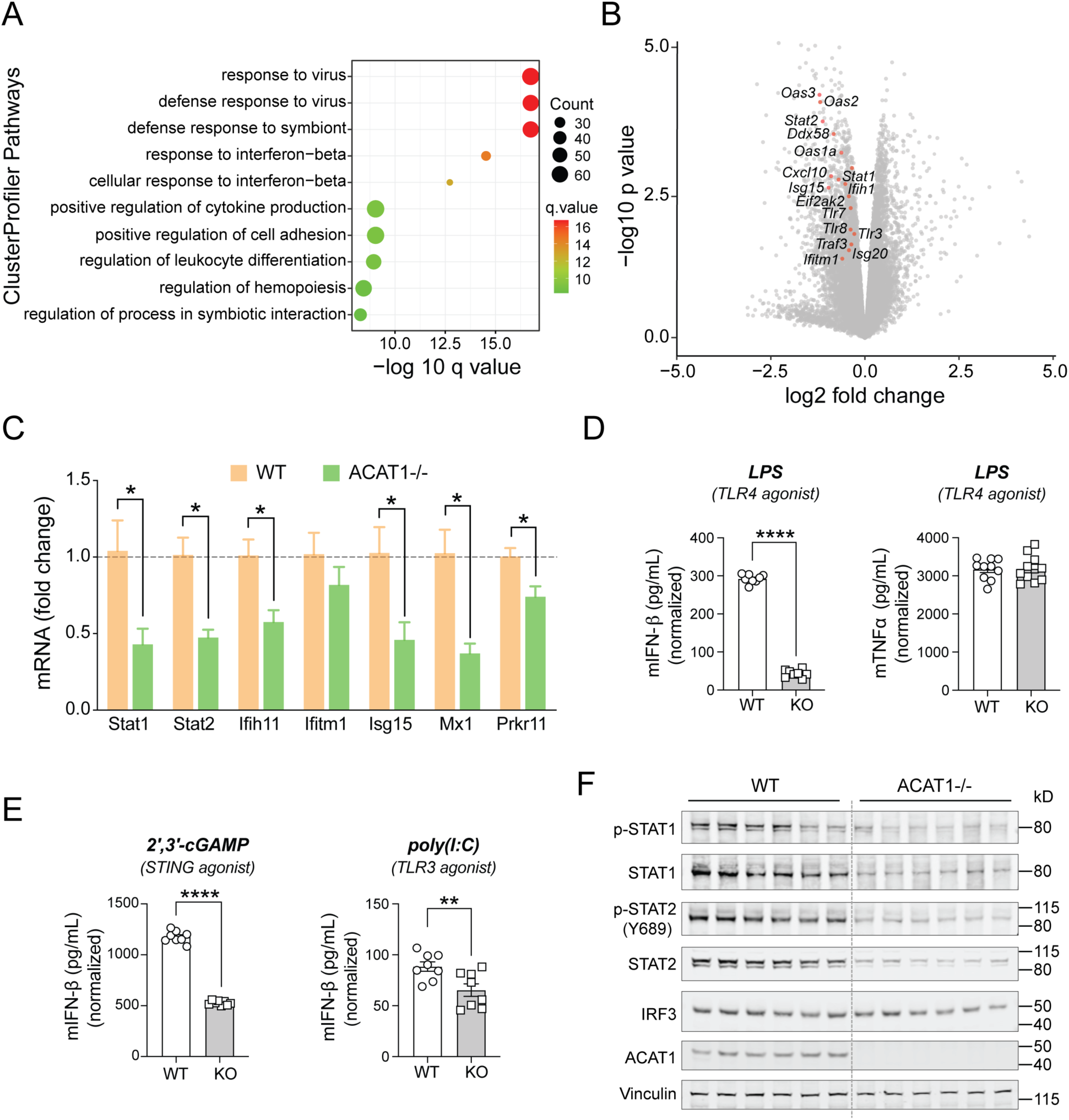
Type 1 Interferon pathway is downregulated in ACAT1 KO macrophage cells. (A) Dot plot of cluster profiler pathway from 1618 downregulated genes out of a total of 3288 common DE genes. The x axis represents negative log10-transformed q values, and the dot-plot color was scaled to transformed q values. The size of the dot was scaled to number of genes overlapping with the indicated pathway on the y axis. (B) Volcano plot of DE genes (p<0.05) from WT vs. ACAT1 KO macrophage cells under basal state (n=3 different lines/genotype). The statistically significant type I interferon genes are highlighted in red circles and labeled.(C) Quantitative RT-PCR analysis of type 1 interferon pathway related genes in WT vs ACAT1 KO cells (n=3 lines/genotype). Data were normalized to 18S rRNA and represented as mean ± SEM. *p<0.05. Unpaired two-tailed Student’s t-test. (D) IFNβ or TNFα production measured by ELISA from WT vs ACAT1 KO cells stimulated with TLR4 agonist. (n=8 replicates/treatment). (E) IFNβ production measured by ELISA from WT vs ACAT1 KO cells stimulated with STING or TLR3 agonist. (n=8 replicates/treatment). (E) Representative immunoblot of phospho-STAT1/2 and total STAT1/2 levels in WT vs ACAT1 KO cells (n=6/genotype).

### ACAT1 Depletion Attenuates Histone Acetylation

Cluster profiler enrichment analysis curated for intracellular signaling pathways also uncovered that chromatin and histone modification pathways were differentially regulated in ACAT1 KO cells comparing to WT counterparts upon LPS stimulation (Figure 2A, and Suppl. Table 2). To evaluate if histone acetylation was altered in the absence of ACAT1, we employed a cellular mitogenic stimulation protocol to induce a net increase in histone acetylation where cells are synchronized through serum starvation and reintroduction (27) (shown in the inset of Figure 2B). Immunoblot analysis in response to serum stimulation over an 8-hour period shows increased acetylation of nuclear H3 and H4 in control cells but not in ACAT1 KO cells (Figure 2B). Interestingly, the extent and dynamic increase in tubulin acetylation in the cytosol appeared similar between WT and ACAT1 KO cells (Figure 2B). The dynamic changes in the degree of H3 acetylation compared to that of tubulin over a 24 hours post-serum stimulation period is shown in (Figure 2C). ACAT1 is a distal enzyme leading to the production of Ac-CoA during FAO and the abundance of acetyl-CoA positively regulates histone acetylation to facilitate gene transcription (27, 28). We therefore measured levels of acetyl-CoA and CoA in WT and ACAT1 KO cells by HPLC. As expected, the levels of acetyl-CoA and CoA were significantly lower in the ACAT1 KO cells (Figure 2D). Besides ACAT1 KO murine macrophage cell line, we also utilized primary human monocytes with siRNA knockdown to examine how ACAT1 regulates histone acetylation through different metabolic pathways using various upstream substrates, e.g. glucose, and the medium chain fatty acid octanoate (Figure 2E and Suppl. Figure 3). Interestingly, octanoate supplementation had the most robust effect on H3 and H4 acetylation and these modifications were blunted following the knockdown of ACAT1 in human monocytes (Figure 2, E and F). In contrast, glucose-mediated histone aceylation in human monocytes was not affected by ACAT1 knockdown (Suppl. Figure 3). To further demonstrate the pivotal role of the fatty acid oxidation pathway on histone acetylation, using siRNA knockdown, we disrupted two key enzymes involved in medium-chain FAO: Acyl-CoA Synthetase Medium Chain Family Member 2A (ACSM2A) and Acyl-CoA Dehydrogenase (ACADM) (Suppl. Figure 4A). ACSM2A knockdown significantly blunted H3 and H4 acetylation following octanoate supplementation (Suppl. Figure 4, B and C). ACADM knockdown had no impact, possibly due to its low expression in monocytes under basal state (data not shown). In parallel, the overexpression of ACAT1 in the presence of octanoate increased the acetylation of histone H3, but not that of the cytosolic cytoskeletal protein α-tubulin (Figure 2, G and H). Furthermore, to determine whether histone acetylation *via* FA derived acetyl-CoA is substantially impaired in ACAT1 KO cells, we used stable isotope tracing of uniformly labeled ^13^C-octanoate incorporation into whole-cell acetyl-CoA pools followed by LC-MS detection of histone H3 K9 and K14 acetylation (Figure 3A). The degree of ^13^C incorporation in acetyl-H3 is markedly reduced in ACAT1 KO macrophage cells (Figure 3B). Overall, these findings provided evidence to support the link between mitochondrial ACAT1 and nuclear histone acetylation through lipid-derived acetyl-CoA.

**Figure 2.**
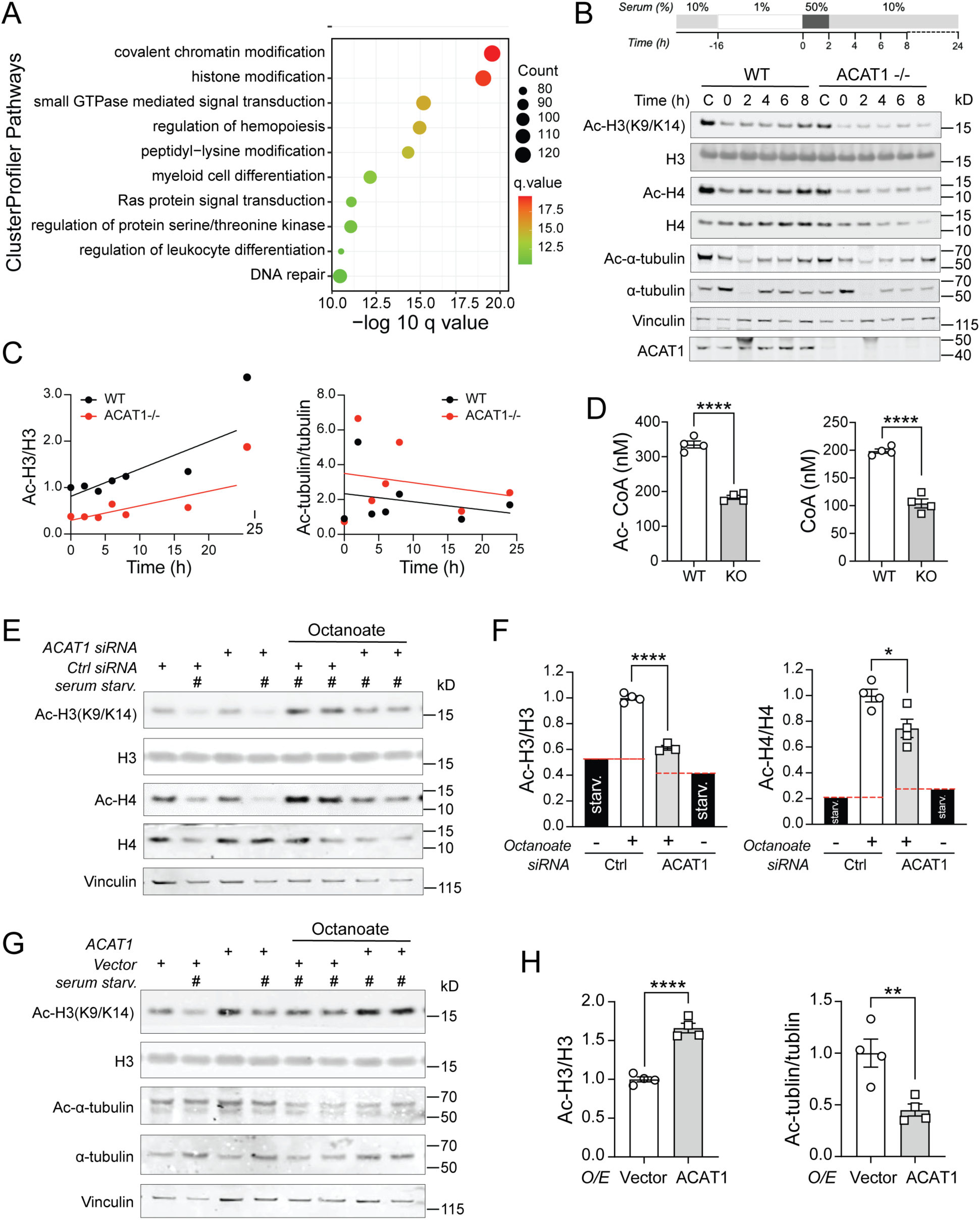
Fatty acids mediated histone acetylation is impaired in ACAT1 deficient cells. (A) Dot plot of cluster profiler pathway from 2760 downregulated genes out of a total of 4255 common DE genes. The x axis represents negative log10-transformed q values, and the dot-plot color was scaled to transformed q values. The size of the dot was scaled to number of genes overlapping with the indicated pathway on the y axis. (B) Scheme and Western blotting of histone acetylation in WT and ACAT1 KO cells during cellular response to serum stimulation. (C) Ratio of acetylated histone H3 to total H3 or acetylated tubulin to total tubulin was quantitated at each time point and fitted to a linear regression curve. (D) Acetyl-CoA or CoA level was assessed in WT and ACAT1 KO cells by HPLC. (E) Western blotting of histone acetylation in human monocytes transfected with either control siRNA or ACAT1 siRNA under serum starvation in combination with octanoate (2mM). (F) Quantitative analysis of acetylated H3 to total H3 or acetylated H4 to total H4 ratio in above siRNA transfected cells. (G) Western blotting of histone acetylation in human monocytes transfected with either empty vector or human ACAT1-expressing plasmid under serum starvation in combination with octanoate (2mM). (H) Quantitative analysis of acetylated H3 to total H3 or acetylated α-tubulin to total α-tubulin ratio in above transfected cells.. Data were analyzed by upaired two-tailed Student’s *t*-test. All data were represented as mean ± SEM. \**p*<0.05; \*\*\*\**p*<0.0001.

**Figure 3.**
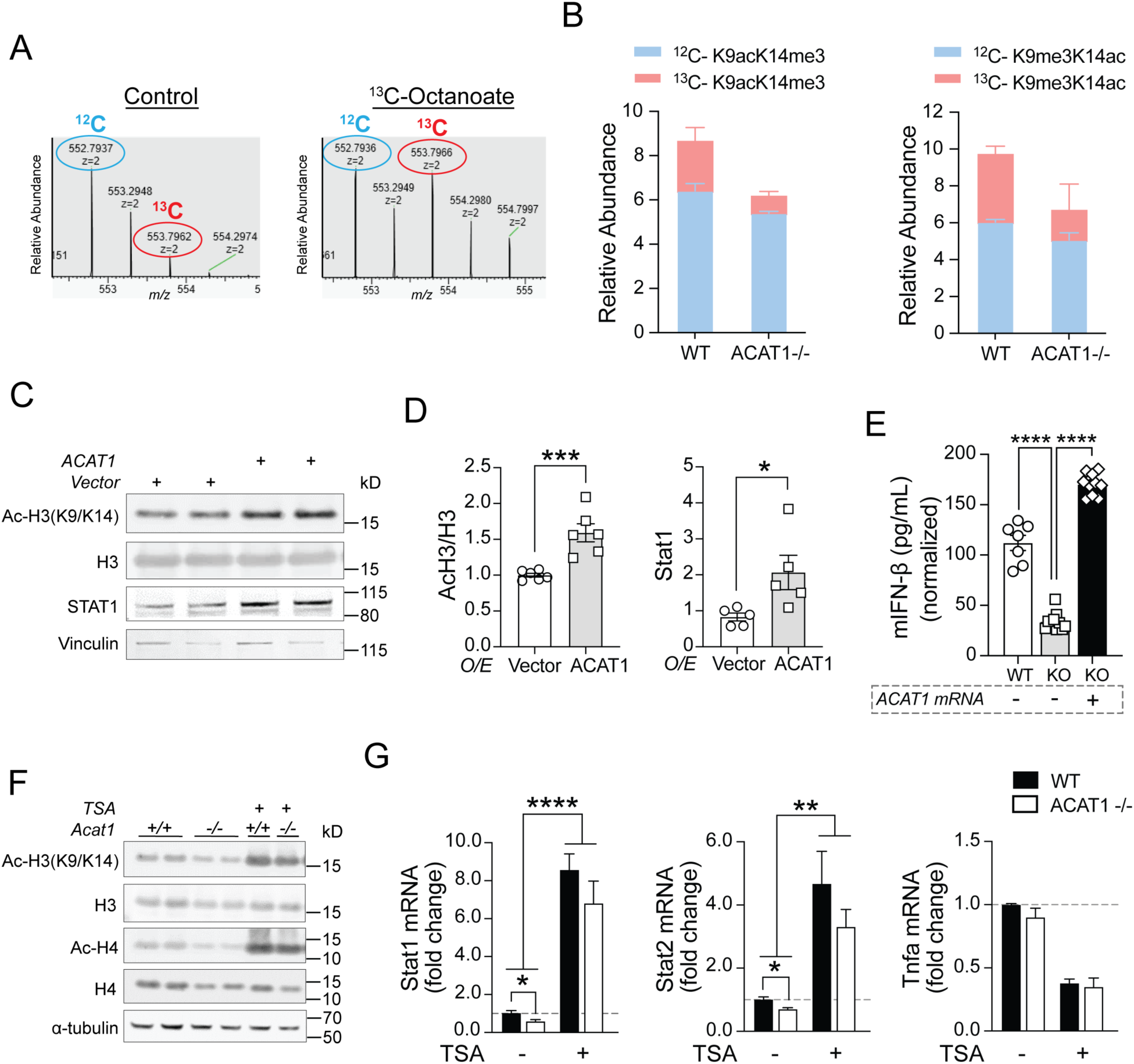
Histone acetylation is required for ACAT1 mediated transcriptional regulation of STAT1 and type 1 interferon production. (A) Representative MS spectra from WT cells treated with vehicle (left) or [U-^13^C]octanoate (right) added to serum-starvation media, highlighting the isotope distribution of histone H3 peptides acetylated on lysines 9 and 14 (z = 2). Mass shifts to the right indicate presence of [U-^13^C]octanoate-derived carbon at those specific histone lysines. (B) Relative enrichments of acetylated ^13^C-K9 or ^13^C-K14 for histone H3 corrected for natural abundance (^12^C) ± SEM. (C) Western blotting of histone H3 acetylation and STAT1 in human monocytes transfected with either vector or ACAT1 under normal serum condition. (D) Quantitative analysis of acetylated H3 to total H3 ratio or STAT1 level in above transfected cells. (E) mouse IFNβ production measured by ELISA from WT, ACAT1 KO cells or KO cells transfected with ACAT1 mRNAs. (F) Western blotting of histone acetylation in WT and ACAT1 KO cells treated with vehicle or TSA. (G) Quantitative RT-PCR analysis of mouse Stat1, Stat2 and Tnfα genes in WT and ACAT1 KO cells treated with either vehicle or TSA for 16hr. Data were normalized to 18S rRNA and represented as mean ± SEM. *p<0.05, **p<0.01, ***p<0.001, ****p<0.0001. Unpaired two-tailed Student’s *t*-test.

### Histone Acetylation is Required for ACAT1 Mediated Upregulation of STAT1 and T1IFN

Histone acetylation, a posttranslational modification known for enabling dissociation of the interaction between histones and DNA to modify nucleosomal conformation, is crucial for facilitating gene transcription. We therefore examined the impact of ACAT1 overexpression on histone acetylation and STAT1 levels in human monocytes. Here ACAT1 overexpression increased H3(K9/K14) acetylation and concominantly increased steady-state levels of STAT1 (Figure 3, C and D). Additionally, when Acat1 was reintroduced into ACAT1 KO J774A.1 cells, there was a concurrent induction of LPS-mediated IFN-β secretion, reaching levels comparable to those observed in WT cells (Figure 3E). To validate the role of histone acetylation in this T1IFN biology, we then employed trichostatin A (TSA), a broad histone deacetylase inhibitor. Here, TSA treatment dramatically augmented histone H3 and H4 acetylation in both WT and KO macrophages without affecting total protein levels (Figure 3F). In parallel, TSA increased the transcript levels of Stat1 and Stat2, but not that of Tnfα in either WT or ACAT1 KO cells (Figure 3G).

Before delving deeper, we employed an integrated bioinformatic approach comparing downregulated DE genes in our RNAseq dataset (WT *vs.* ACAT1 KO) with a public H3K9ac CHIPseq dataset from wildtype murine bone marrow-derived macrophages (GSE113226). The Venn diagram illustrates an overlap of 637 genes between these datasets (Figure 4A). Subsequent ClusterProfiler pathway analysis supported a significant overlap of genes associated with the antiviral response in these datasets (Figure 4B). To examine potential modifications in the transcriptional machinery of Stat1 in the presence or absence of ACAT1, we used the UCSC genome browser (28) to annotate the promoter and enhancer loci of Stat1 (Figure 4C). Quantitative H3K9ac CHIP-murine Stat1-PCR revealed that the deletion of Acat1 resulted in diminished H3K9ac binding at the putative promoter and selective distal enhancers of the murine Stat1 gene (Figure 4D). These findings suggested that ACAT1 could influence Stat1 gene expression through histone acetylation and its binding to the upstream regulatory elements of the Stat1 locus.

**Figure 4.**
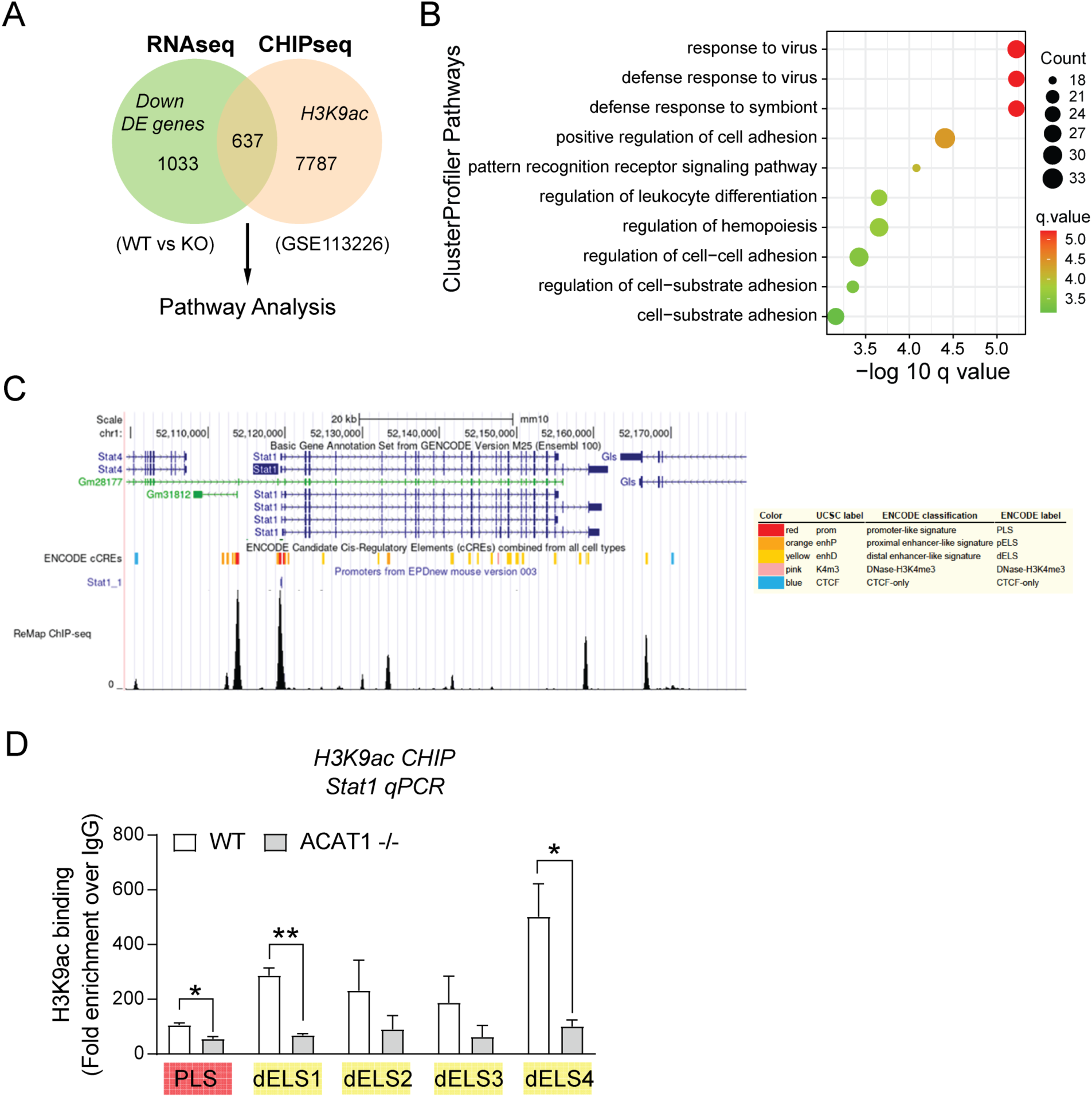
Binding of acetylated histone H3 to Stat1 promoter regions are reduced in ACAT1 KO cells. (A) Flowchart showing downregulated DE genes from RNAseq dataset and H3K9ac CHIPseq dataset (GSE113226) used to identify DE genes specifically regulated by ACAT1 and H3K9 acetylation. The 637 overlapping DE genes were subjected to pathway enrichment analysis. (B) Dot plot of cluster profiler pathway from above 637 overlapping DE genes out of a total of 2969 common DE genes. The x axis represents negative log10-transformed q values, and the dot-plot color was scaled to transformed q values. The size of the dot was scaled to number of genes overlapping with the indicated pathway on the y axis. (C) UCSC Genome Browser on mouse Stat1 locus. (D) Quantitative H3K9ac CHIP-PCR analysis of mouse Stat1 promoter/enhancer sequences in WT and ACAT1 KO cells. Data were represented as mean ± SEM. *p<0.05, **p<0.01. Unpaired two-tailed Student’s t-test.

### The Carnitine Shuttle is Required for ACAT1 Mediated Acetylation of Histone H3

Acetyl-CoA cannot directly traverse organelle membranes, and thus, a transport mechanism across the mitochondrial membranes is required for the FAO mediated nuclear-cytosolic acetylation events. The pathways orchestrating histone acetylation from mitochondrial derived acetyl-CoA have been shown to be dependent (27) or independent on the ATP-citrate lyase (ACLY) (26, 29). In the latter case, recent study indicates both glucose and fatty acids can supply nuclear-cytosolic acetyl-CoA through a pathway that does not require ACLY but acetylcarnitine shuttling (27, 29). We sought to understand which cellular mechanism is involved in the ACAT1 mediated histone acetylation from FA-derived carbon. Unlike ACAT1, the siRNA knockdown of ACLY did not disrupt histone H3 and H4 acetylation in human monocytes after serum-starvation overnight, followed by treatment with octanoate (Figure 5A). We then disrupted various components of acetylcarnitine shuttling including carnitine acetyltransferase (CrAT) and the carnitine/acylcarnitine translocase (CACT). Interestingly, following overexpression of ACAT1 in human monocytes, the siRNA knockdown of CrAT to a lesser extent, and the knockdown of CACT to a greater extent, blunted histone H3 acetylation without altering the degree of α-tubulin acetylation (Figure 5, B and C). These data support that acetyl-CoA generated through mitochondrial ACAT1 is dependent on the acetylcarnitine shuttle for histone acetylation.

**Figure 5.**
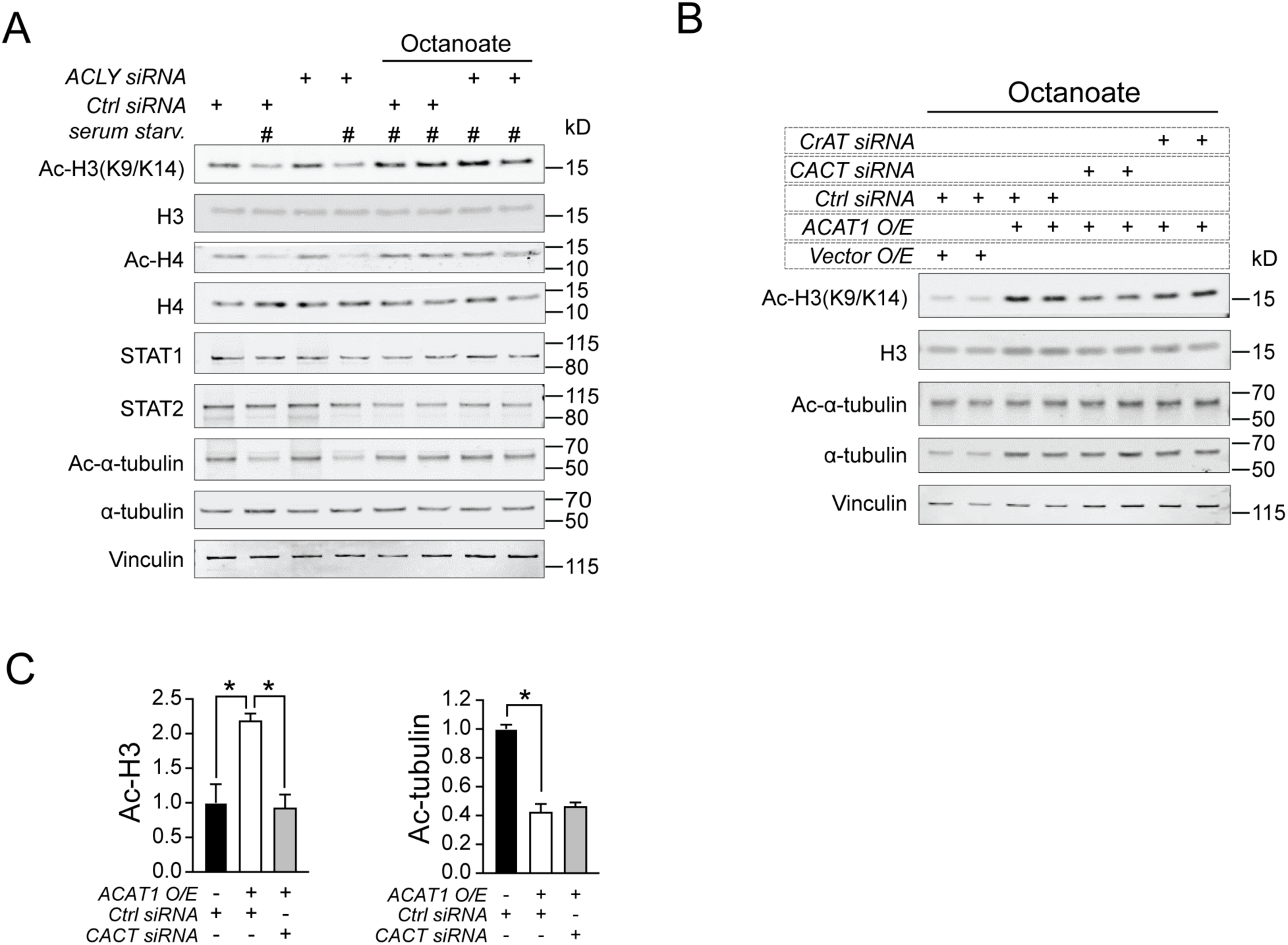
Carnitine shuttle is required for the action of ACAT1 on histone acetylation. (A) Western blotting of histone acetylation, STAT1 and STAT2 in human monocytes transfected with either control siRNA or ACLY siRNA under serum starvation in combination with octanoate (2mM). (B) Western blotting of histone and tubulin acetylation in human monocytes transfected with either empty vector and control siRNA or ACAT1 and/or control siRNA, CACT siRNA, CrAT siRNA under serum starvation in combination with octanoate (2mM). (C) Quantitative analysis of acetylated H3 to total H3 or acetylated α-tubulin to total α-tubulin ratio in above transfected cells. Data were represented as mean ± SEM. *p<0.05. Unpaired two-tailed Student’s *t*-test.

### ACAT1 and histone acetylation are induced in monocytes from Obese compared to Lean human subjects

As obesity is associated with disrupted lipid metabolism and low-level inflammation, we sought to investigate whether the T1IFN pathway modulated by ACAT1 is constitutively activated in individuals with obesity. To explore this, we utilized integrative bioinformatics comparing DE genes between WT *vs.* ACAT1 KO murine macrophage, translated into a human database (30), with a RNAseq dataset comparing PBMC gene expression in lean *vs*. obese subjects (31). The analysis identified DE genes blunted in ACAT1 KO cells and induced in obese PBMCs, revealing an overlap of 210 genes (Figure 6A). Pathway analysis of these overlapping DE genes highlighted that the most significantly divergent regulation involved genes associated with type 1 IFN signaling (Figure 6B). In a cohort of lean and obese, otherwise healthy subjects, recruited under our human Disease Discovery Protocol (NCT01143454), research blood was collected to perform primary monocytes’ isolation using negative selection. qRT-PCR from these monocytes revealed evident induction of transcripts related to T1IFN pathway in obese subjects, including STAT1, CXCL10, ISG15, MX1 and SOCS3 (Figure 6C). Immunoblot analysis on monocytes from the same subjects showed that obesity was associated with increased ACAT1 expression and elevated histone H3 and H4 acetylation (Figure 6D). Collectively, these findings suggest that ACAT1-mediated histone acetylation on the transcriptional regulation of type 1 interferon pathway could be dysregulated in patients with obesity.

**Figure 6.**
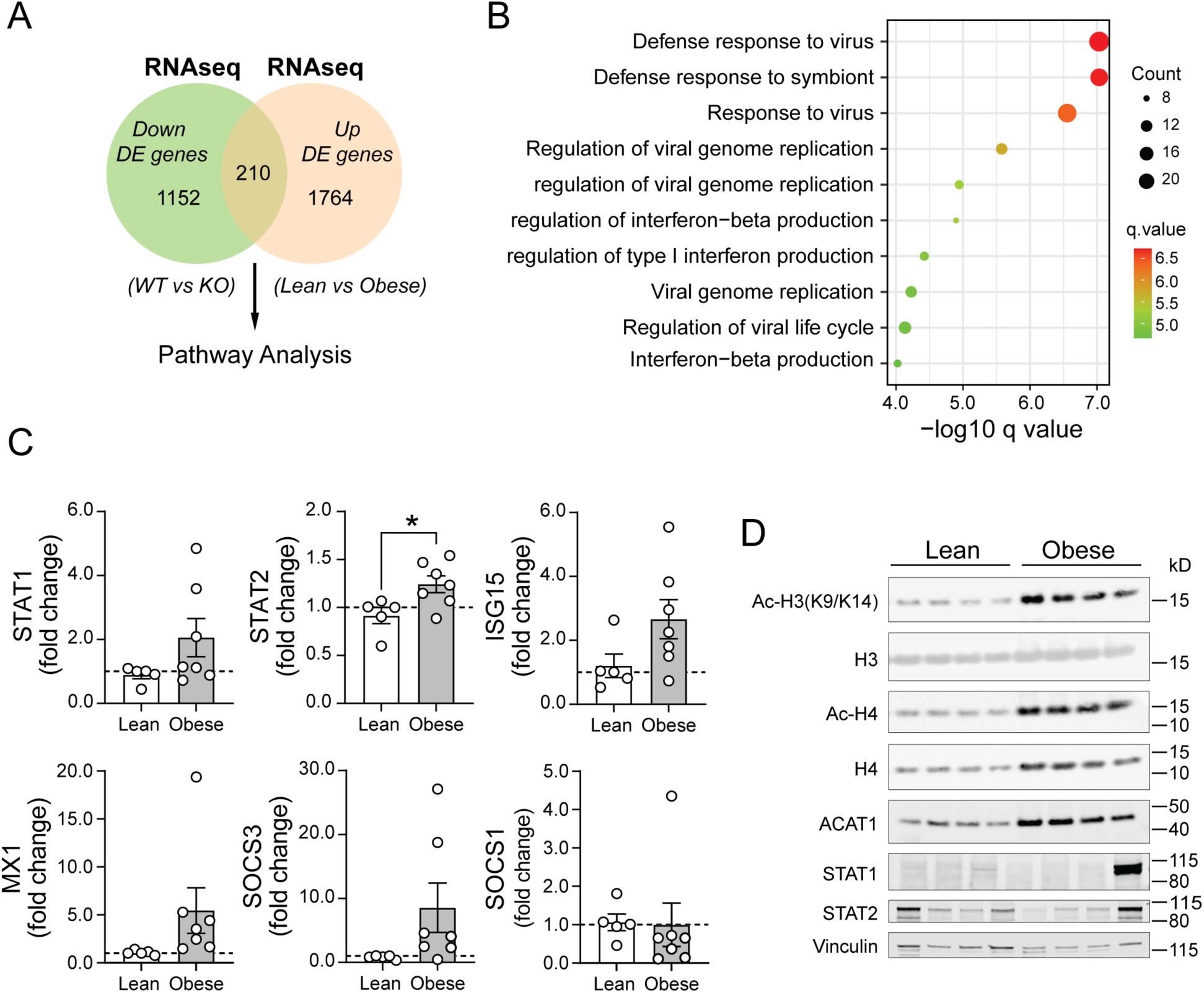
ACAT1 and histone acetylation levels are elevated in human subjects with obesity. (A) Flowchart showing an overlap of 210 genes from downregulated DE genes in ACAT1 KO RNAseq dataset and upregulated DE genes from Obese PBMC RNAseq dataset used to identify pathways specifically regulated by ACAT1 and obesity. (B) Dot plot of pathway analysis from above 210 overlapping DE genes out of a total of 2969 common DE genes. The x axis represents negative log10-transformed q values, and the dot-plot color was scaled to transformed q values. The size of the dot was scaled to number of genes overlapping with the indicated pathway on the y axis. (C) Quantitative RT-PCR analysis of type 1 interferon pathway related genes in monocytes from lean and obese subjects. (n=4-5 subjects/group). Data were normalized to 18S rRNA and represented as mean ± SEM. *p<0.05. Unpaired two-tailed Student’s t-test. (D) Immunoblot of histone acetylation and ACAT1 in lean *vs* obese monocytes. (n=4 subjects/lean and obese group).

## DISCUSSION

In this study we uncover that acetyl-CoA generated by octanoate oxidation plays an important regulatory role in the induction of T1IFN signaling in murine and human macrophages. This effect is dependent on mitochondrial ACAT1, the carnitine/acylcarnitine translocase and histone acetylation. This histone acetylation in turn, upregulates the canonical T1IFN signaling and transcriptional regulatory mediator STAT1. Furthermore, preliminary data implicate that this program may be induced and operational as a component of low-grade inflammation in human subjects with obesity (32).

The concept that metabolic pathways can play an important role in innate and adaptive immunity is now well established and described as immunometabolic regulation. At the level of individual metabolites functioning as signaling intermediates, evidence supports immunomodulatory roles TCA cycle, purine pathway and amino acid metabolites, as well as short chain fatty acids and ketones (3, 11, 14, 33, 34). In this context, acetyl-CoA is emerging as an immunomodulatory metabolite in its role as a mediator of histone acetylation downstream of glucose and fat catabolism (27, 29). Glucose induced histone acetylation is orchestrated by its oxidative metabolism to generate citrate in the TCA, citrate transported to the cytosol and to the nucleus, where it is converted by ATP-citrate lyase into acetyl-CoA, for subsequent histone acetylation. In the context of immunometabolism, TLR4 activation via MyD88 and TRIF increases glucose oxidation-dependent, ACLY-mediated histone acetylation to promote LPS-inducible gene sets in murine macrophages (35). Furthermore, in a human macrophage transformed cell line, cGAS-STING and TBK1 driven cytokine production is similarly dependent on glucose oxidation and ACLY levels (36). Interestingly, in hepatocytes ^13^C-tracing studies show that the oxidation of the medium fatty acid octanoate becomes the preferential substrate for histone acetylation at the expense of glucose and glutamine catabolism (26). Furthermore, that study showed that octanoate-driven histone acetylation was independent of ACLY (26). Recently, acetylcarnitine shuttling has been implicated as a mechanism whereby two-carbon units can be transferred from the mitochondria to the cytosol, to support histone acetylation (29). In this study we further define this pathway, by showing that in macrophages, carnitine/acylcarnitine translocase (CACT), and to a lesser extent carnitine acetyltransferase (CrAT), are required for this octanoate directed histone acetylation. Moreover, we show that this system is operational in the induction of T1IFN in macrophages and that this process is dependent on ACAT1 in the FAO pathway.

The role of lipids in immunometabolism has predominantly been evaluated in the context of arachidonic acid and prostaglandins effects (37–39) and by immunomodulatory roles of specific lipid species including saturated (40), and short chain fatty acids (11). Additionally, FAO itself has been shown to be necessary for the integrity and function of CD8^+^ memory T cells (41) and for CD4^+^ T regulatory cells (42). This study expands on this work to show the importance of FAO in macrophages to control inflammatory gene histone acetylation, and in this instance, via the regulation of STAT1 for the subsequent transactivation of T1IFN signaling.

Interestingly, ACAT1 has pleiotropic ‘substrates’ with its’ role in the catabolism of fatty acids, isoleucine, and ketone bodies. Furthermore, mitochondrial ACAT1 also facilitates the acetylation of components of the pyruvate dehydrogenase complex to modulate glucose homeostasis (22). In that study, ACAT1-mediated acetylation of components of this complex inhibit its activity to promote glycolysis over glucose oxidation, to functionally promote tumor cell proliferation (22). Interestingly, although not established, this mechanism could possibly also be operational in the preferential use of fatty acids for histone acetylation in the presence of octanoate.

In conclusion, this study contributes to the emerging body of evidence that various aspects lipid metabolism and different lipid metabolites contribute to the expananding knowledge base in the role of metabolic pathways and metabolites in the control of immune function. Furthermore, this study uncovers that the role of FAO may play an important epigenetic role in myeloid cells in the control of type I interferon signaling. At the same time our initial data comparing monocytes in lean and obese individuals implicate that this program may also be operational in nutrient-level linked immunomodulation in human disease pathophysiology.

## MATERIALS AND METHODS

### Generation of ACAT1 KO murine macrophage J774A.1 cell line

WT and ACAT1 KO cells were generated by transfecting a pSpCas9-2A-GFP vector with no guide RNA or containing a guide RNA targeting the fourth exon of murine Acat1 gene. Following transfection, cells were sorted by FACS based on GFP fluorescence and plated to 96-well plate at a density of ∼5 cells per well (not all sorted cells may survive the FACS process or the subsequent outgrowth; therefore, the number of cells sorted was adjusted to five instead of one). Single-cell clonal populations for both WT and ACAT1 KO were generated and expanded for 2-3 weeks. The indel mutation at the target loci was verified by genomic DNA sequencing. The protocol was followed as previously published (43).

### RNA Sequencing and Bioinformatics Analysis

Total RNA was extracted with the NucleoSpin® RNA kit (Takara) and RNA quality was assessed by Agilent Bioanalyzer. Libraries were prepared using TruSeq stranded mRNA kit (Illumina) and sequenced in a HISeq 3000 (Illumina) by the DNA Sequencing and Genomics Core at NHLBI. FastQC (http://www.bioinformatics.babraham.ac.uk/projects/fastqc) was used to confirm quality of RNA seq fasta files. Adaptor trimming, RNA sequence alignment, PCA analysis, differential pathway analysis was performed as previously described (20). Genes with *p* value <0.05 were considered DE genes.

### Human Monocyte Cultures

The National Heart, Lung, and Blood Institute (NHLBI) Institutional Review Board (IRB) approved this study (NCT01143454). Lean and obese otherwise healthy volunteers signed informed consent before donating blood for analysis. Primary peripheral blood mononuclear cells (PBMCs) were isolated from human blood by density centrifugation using Lymphocyte Separation Medium (MP Biomedicals). Human monocytes were negatively selected from PBMCs using the monocytes Isolation Kit (Miltenyi Biotec). Monocytes were then plated 0.15×10^6^/well onto 96-well plate (for ELISA) or 10^6^/well onto 12-well plates (for RNA isolation or Western blotting) in RPMI media supplemented with 10% human serum. Human whole blood and elutriated monocytes were obtained from blood bank of NIH.

### Inhibitors, siRNA and Nucleofection of human monocytes

HDAC inhibitor Tricostatin A (TSA) (Cayman) was used in cell culture at 50µM for 16hr before subsequent assays. ON-TARGET*plus* siRNA for knocking down gene expression of ACAT1, ACLY, CRAT, CACT or non-targeting control siRNA was purchased from Horizon Discovery. siRNA against human genes or control siRNA were incubated with a mixture of nucleofection solution (Human Monocytes Nucleofector kit; Lonza) and primary human monocytes and placed in nucleofection cuvettes subjected to program Y-010 for the Nucleofector 2b Device (Lonza). 500µL RPMI medium was immediately added into cuvettes after nucleofections. Cells were then plated in 12-well plate and incubated at 37°C under 5% CO_2_ for 48hr before harvesting for the assays. Plasmid transfections using empty vector (pcDNA3.1) or human ACAT1-expression constructs were conducted with the same protocol as siRNA transfection.

### RNA Isolation and Quantitative PCR analysis

Total RNA was extracted from monocytes using NucleoSpin® RNA kit (Takara) and RNA concentration was measured using NanoDrop Spectrophotometer (Thermo Fisher Scientific). cDNA was synthesized with the SuperScript™ III First-Strand Synthesis System for RT-PCR (Thermo Fisher Scientific) according to the manufacturer’s instructions. Quantitative real-time PCR was performed using FastStart Universal SYBR Green Master (Rox) (Roche Holding) and run on LightCycler 96 Systems (Roche Holding). Transcript levels of STAT1, STAT2 and 18s rRNA were measured using validated gene-specific primers (QIAGEN). Primers for ACAT1, ACLY, CRAT, CACT, ACSM2A and ISGs were custom synthesized at IDT Inc. Relative gene expression was quantified by normalizing cycle threshold values with 18S rRNA using the 2^-ΔΔCt^ cycle threshold method.

### Western Blotting

Human monocytes were lysed using RIPA buffer supplemented with protease inhibitor cocktail (Roche) and phosphatase inhibitors (Sigma-Aldrich). The lysates were separated by NuPAGE™ 4-12% Bis-Tris Protein Gels (Thermo Fisher Scientific) and transferred to nitrocellulose membranes using Trans-Blot® Turbo™ Transfer System (Bio-Rad Laboratories) according to the manufacturer’s instructions. Membranes were blocked with 50% Odyssey Blocking Buffer in PBS-T (0.1% Tween20 in PBS) buffer and incubated with appropriate antibodies overnight at 4°C. Antibodies used included: STAT1, phospho-STAT1(Y701), STAT2, Acetyl-Histone H3 (Lys9/Lys14), Histone H3, Histone H4 (Cell Signaling Technologies); phospho-STAT2 (Y689), Acetyl-Histone H4 (Lys5/Lys8/Lys12/Lys16) (Millipore); IRF3 (Abcam); and ACAT1 and Vinculin (Sigma). Primary antibody incubations were followed by incubation with IRDye® Secondary antibodies for 1hr at room temperature. Immunoblots were visualized and imaged using Odyssey CLx Imaging System (LI-COR Biosciences). Protein band intensity was measured using Image J software and normalized to Vinculin.

### Cell Stimulation and Cytokine Assays

WT and ACAT1 KO J774A.1 cells were incubated at 0.08 × 10^6^ cells per well in a 96-well plate in complete DMEM medium (10% FBS in DMEM medium) stimulated with 100 ng/mL LPS (Ultrapure Salmonella minnesota R595; Enzo Life Sciences), or 2µg/mL 2’,3’-cGAMP (InvivoGen), or 2µg/mL poly(I:C) (InvivoGen) for 16 h. Cell culture supernatants were collected, centrifuged to remove cells and debris, and stored at –80°C for later analysis. Cytokines of mouse IFNβ and TNFα were assayed by ELISA (R&D Systems). Results were normalized to cell number, as determined by the CyQuant cell proliferation assay (Invitrogen).

### Acetyl-CoA and CoA analysis by HPLC

WT and ACAT1 KO cells were plated at a density of 5×10^6^ cells per 10-cm culture dish and cultured in DMEM (10%FBS) for one day. Cells were washed with PBS, detached from the dish using CellStripper Solution. Cell pellet was collected by centrifugation, washed with PBS twice, and resuspended in 100 μl of 5% 5-Sulfosalicylic acid (Sigma) solution. To permeabilize the cells, samples were frozen in liquid nitrogen and thawed on ice. This freeze–thaw cycle was repeated twice. After centrifugation, the supernatant was filtered using Ultrafree-MC LH Centrifugal Filter (Millipore, Billerica, Massachusetts, UFC30LH25). Then, the samples were transferred into SUN-SRi Glass Microsampling Vials (Thermo Fisher Scientific, 14-823-359) with SUN-SRi 11mm Snap Caps (Thermo Fisher Scientific, 14-823-379), and 80 μl of each sample was separated using an Agilent 1100 HPLC (Agilent Technologies, Santa Clara, California) equipped with a reverse phase column, Luna 3 μm C18(2) 100 Å, 50 x4.6mm, 3 μm (Phenomenex, Los Angeles, California). The detailed protocol was described previously (44).

### Stable Isotope Labeling

WT and ACAT1 KO cells were seeded at a density of 6 × 10^6^ cells per 10-cm culture dish in DMEM containing 10% FBS. The following day, the media was changed to DMEM media containing 1% dialyzed FBS or 1% dialyzed FBS supplemented with 2mM [U-^13^C_8_]-octanoate (Cambridge Isotope Laboratories). After 24h, cells were washed with ice-cold Dulbecco’s phosphate buffered saline and collected into tubes. The cells were further washed twice with ice-cold DPBS, and the cell pellet was frozen at −80°C until extraction.

### Histone H3 Acetylation Analysis by LC-MS/MS

Histones were extracted by following the histone extraction protocol (Abcam) for western blots. Briefly, cells were washed with ice cold PBS, scraped and collected. Cell pellets were then resuspended in Triton extraction buffer (TEB) and lysed on ice while rotating at 4 °C for 10 min. Nuclei were then spun down by centrifugation at 6,500xg for 10 mins at 4°C. Supernatant was then discarded and nuclei were washed with TEB buffer and centrifuged at 6,500xg for 10 min at 4°C. Pellet was resuspended in 0.2N HCL and rotated overnight at 4°C to acid extract histones. Samples were then centrifuged as before and supernatant containing histone proteins were frozen at -20°C. Acid-extracted histones (in 0.2N HCl) were neutralized with pH8 Tris, then spiked with enough 8M Urea Lysis Buffer to achieve a final concentration of 1.5 M urea. Samples were sonicated, reduced, and alkylated. Before trypsinization, the histones were chemically derivatized using propionic anhydride. Trypsin was then added at 50:1 (protein:enzyme) and samples were incubated at 37°C overnight and desalted by SPE, and dried down in 95% and 5% aliquots. For each sample, the larger aliquot (95%) was subjected to acetyl IP and the eluate analyzed by LC-MS/MS as described previously (26).

### Statistical Analysis

Statistical analysis was performed using Prism 9 software (GraphPad) and results are represented as mean ± s.e.m. unless otherwise indicated. Comparisons of two groups were calculated using paired or unpaired two-tailed Student’s *t*-test. Comparisons of more than two groups were calculated using one-way analysis of variance (ANOVA) with Sidak multiple comparisons test or Dunnett’s multiple comparisons test. For all tests, *p* < 0.05 was considered significant.

## Acknowledgments

This research was supported by the NHLBI Division of Intramural Research (MNS – ZIA-HL005199). We thank and acknowledge the assistance of the NHLBI DNA Sequencing and Genomics Core in performing the RNA library sequencing. We would like to thank the NCI Protein Characterization Laboratory for conducting the LC-MS/MS on histone acetylation analysis. We also thank and acknowledge the assistance of Russell Stein on monocytes isolation from human subjects blood samples.

