## Supplemental Materials for "The mitochondrial thiolase ACAT1 regulates monocyte/macrophage type I interferon *via* epigenetic control"

**Suppl. Figure 1.** Characteristics of human ACAT1 obtained from public databases.

**Suppl. Figure 2.** PCA plots of the differentially expressed (DE) genes comparing WT vs ACAT1 KO cells.

**Suppl. Figure 3.** Glucose mediated histone acetylation is intact in ACAT1 deficient cells.

**Suppl. Figure 4.** Fatty acids mediated histone acetylation is impaired in ACSM2A deficient cells.

**Suppl. Figure 5.** Validation of various genes' knockdown by qRT-PCR.

****Suppl. Table 1.** Differentially expressed genes comparing WT *vs* ACAT1 KO under basal state.

****Suppl. Table 2.** Differentially expressed genes comparing WT *vs* ACAT1 KO upon LPS stimulation.

**Above Suppl. Tables can be accessed through the following link to Figshare:
https://figshare.com/s/35cab2ec708b2132d3f1

#### Supplementary Figure 1

A

**ACAT1 expression**

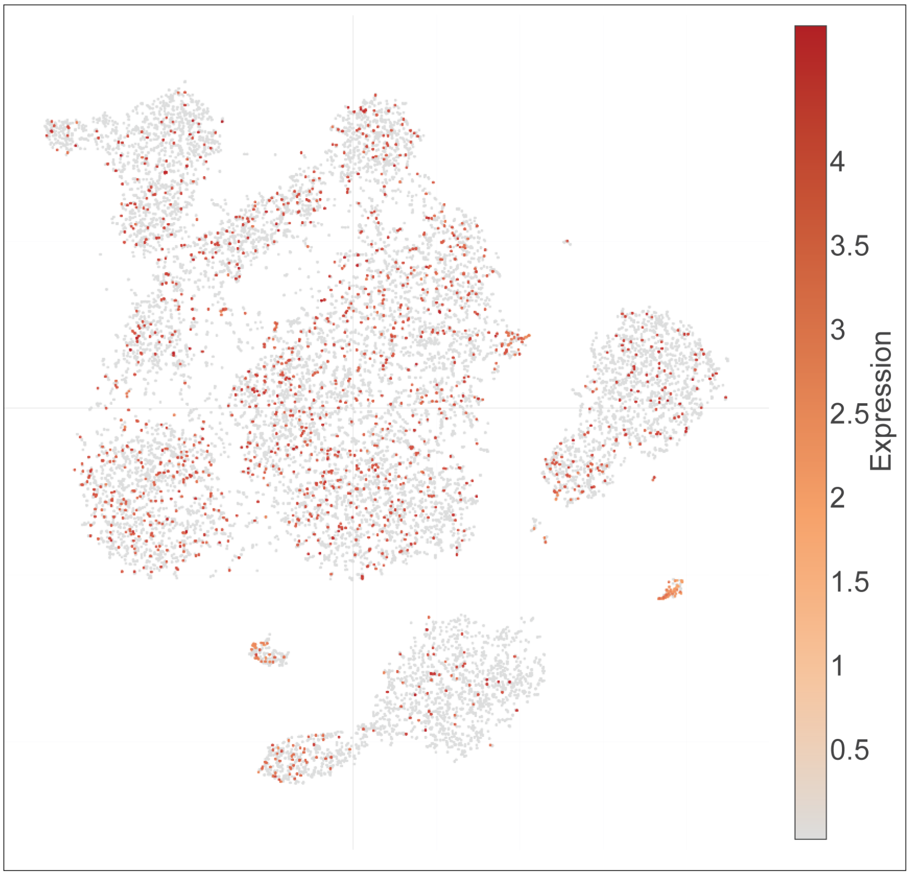


lineage_kmeans

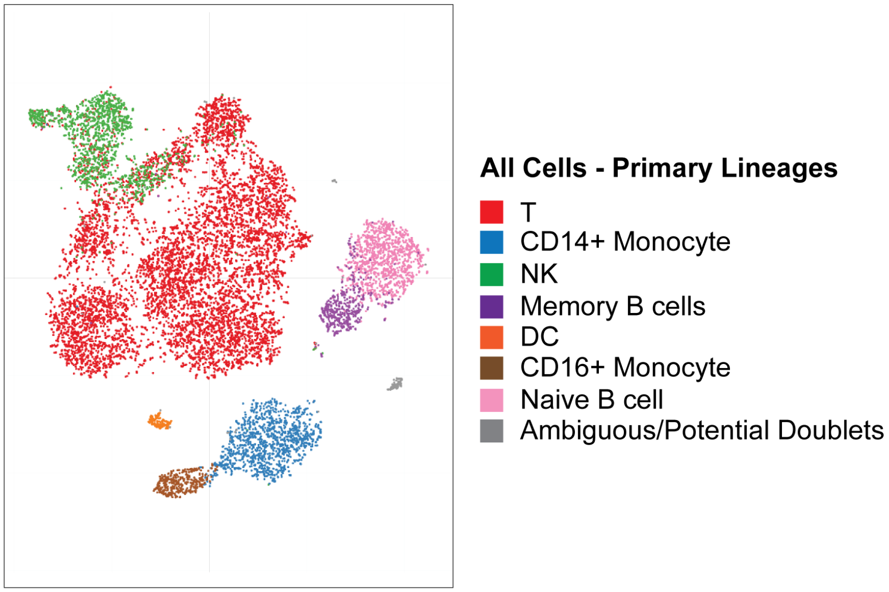


lineage_kmeans

#### Supplementary Figure 1

B

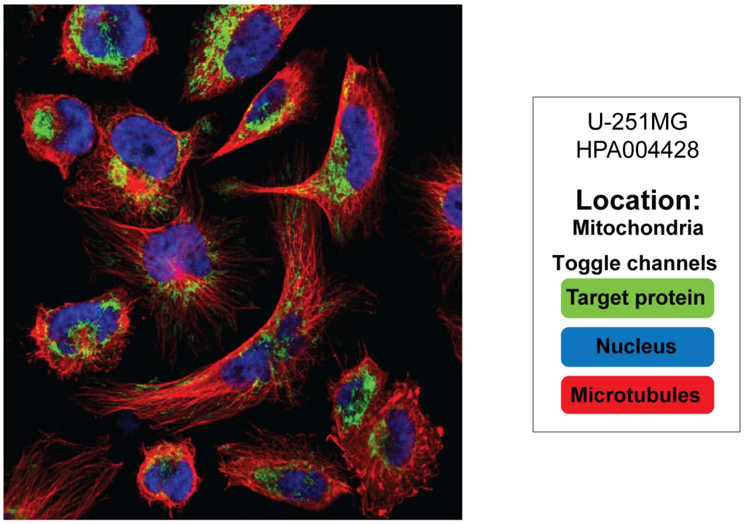


**Figure S1. Characteristics of human ACAT1 obtained from public databases.**

(A) Human ACAT1 expression derived from human PBMC single cell RNA-seq datasets. The resource link is: https://www.immgen.org/ (B) Human ACAT1 cellular localization derived from Human Protein Atlas project. The resource link is: https://www.proteinatlas.org/.

#### Supplementary Figure 2

A

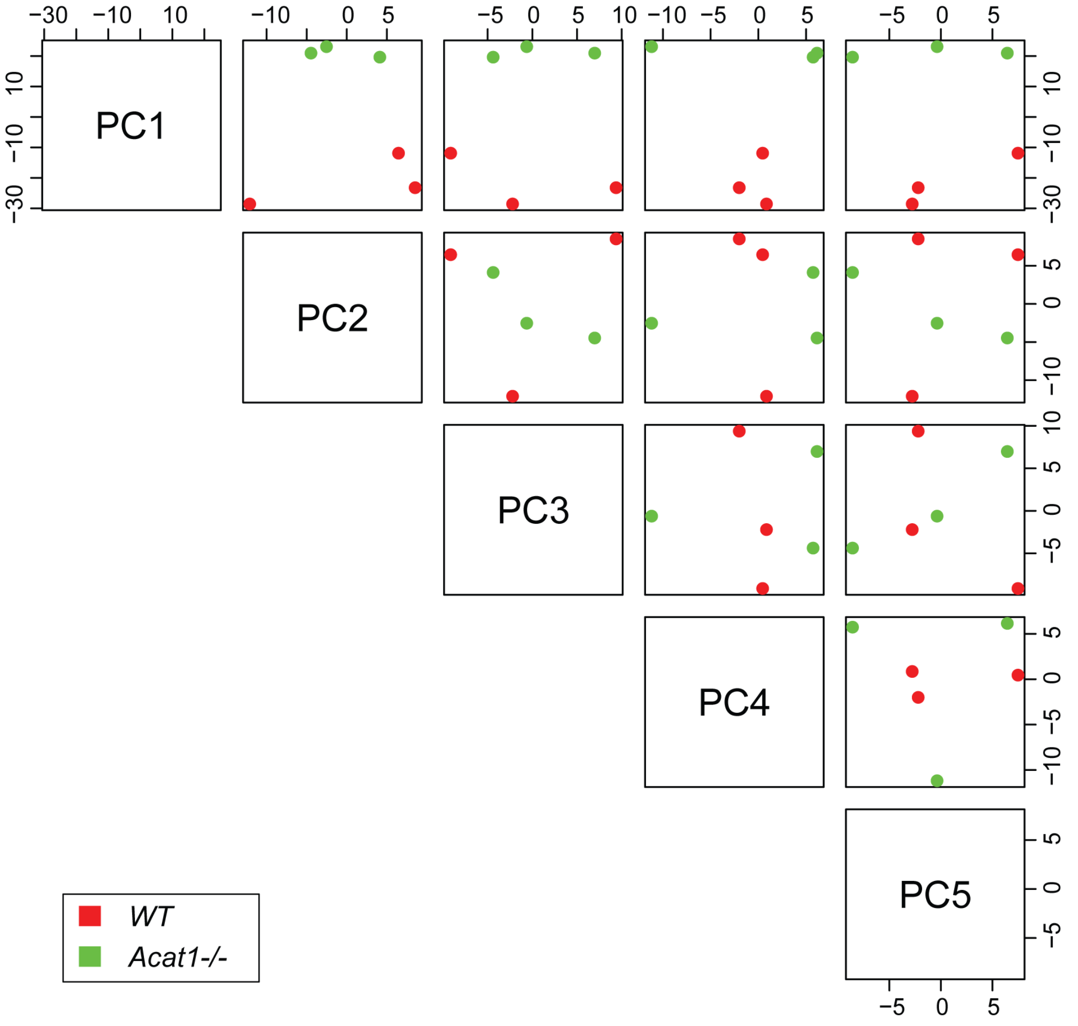

#### Supplementary Figure 2

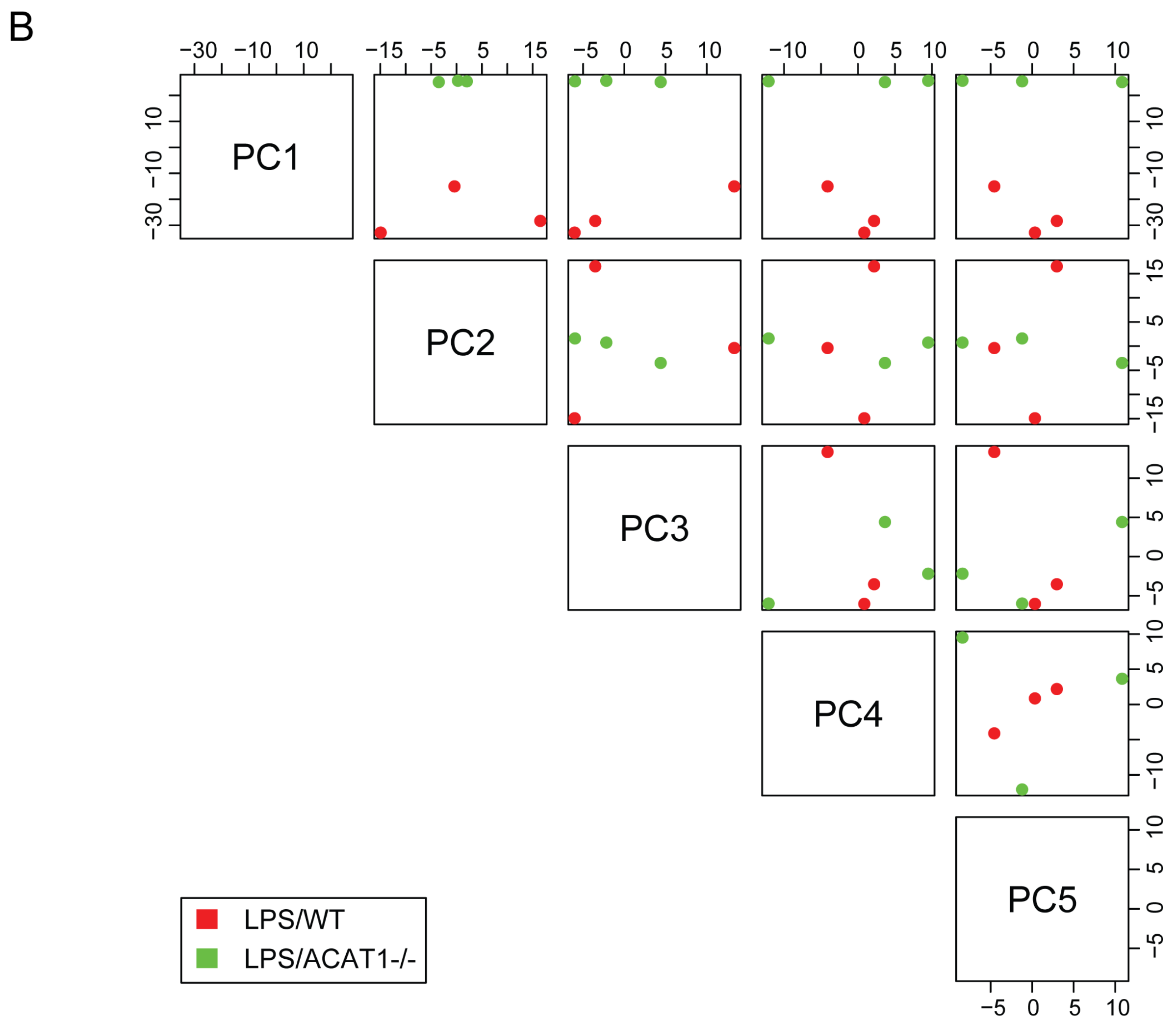


**Figure S2. PCA plots of the differentially expressed (DE) genes comparing WT *vs* ACAT1 KO cells.**

(A) PCA plots of DE genes from WT *vs* ACAT1 KO under basal state. (B) PCA plots of DE genes from WT *vs* ACAT1 KO upon LPS stimulation.

#### Supplementary Figure 3

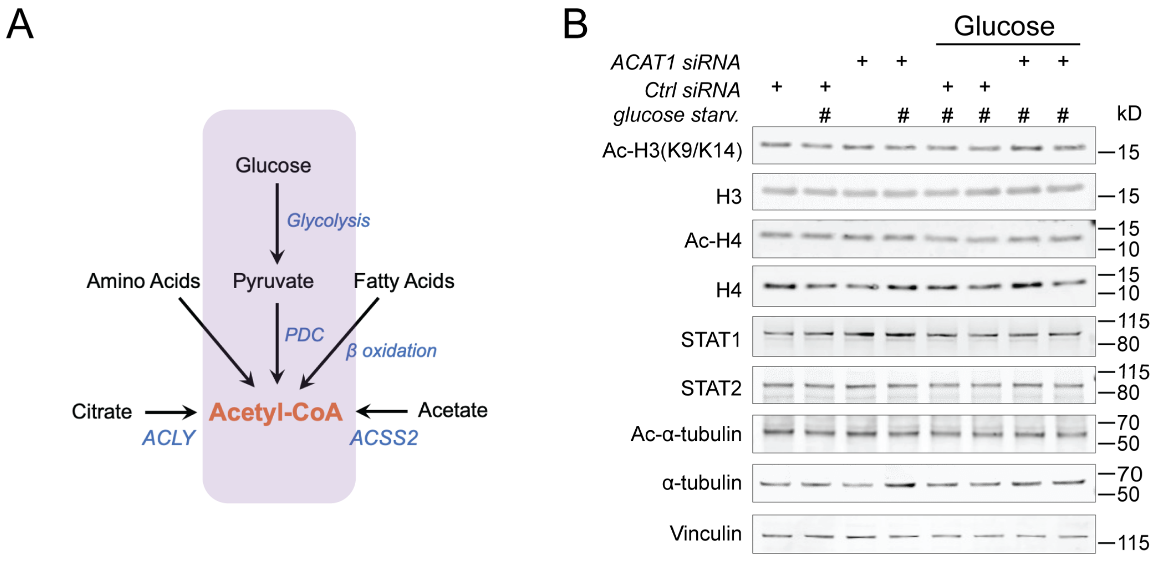


**Figure S3. Glucose mediated histone acetylation is intact in ACAT1 deficient cells.**

(A) Scheme summarizing the major metabolic pathways to generate acetyl-CoA with glucose oxidation pathway highlighted in light purple. (B) Western blotting of histone acetylation in human monocytes transfected with either control siRNA or ACAT1 siRNA under glucose starvation in combination with glucose (11.11mM).

### Supplementary Figure 4

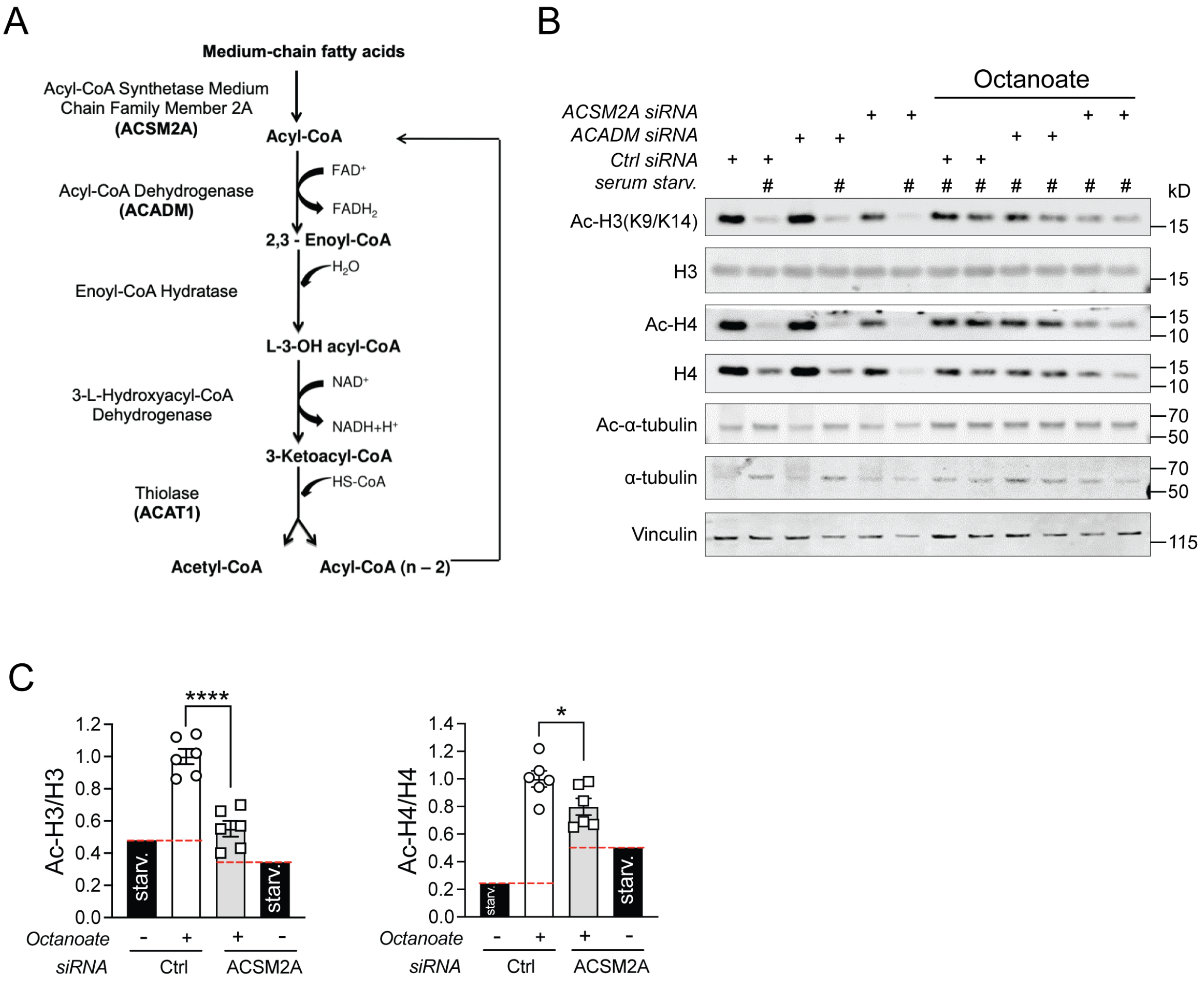


**Figure S4. Fatty acids mediated histone acetylation is impaired in ACSM2A deficient cells.**

(A) Diagram showing medium-chain fatty acids β-oxidation pathway and key enzymes. (B) Western blotting of histone acetylation in human monocytes transfected with either control siRNA, ACADM siRNA or ACSM2A siRNA under serum starvation in combination with octanoate (2mM). (C) Quantitative analysis of acetylated H3 to total H3 or acetylated H4 to total H4 ratio in control siRNA or ACSM2A siRNA transfected cells. Data were analyzed by unpaired two-tailed Student's t-test. All data were represented as mean ± SEM. *p<0.05; ****p<0.0001.

#### Supplementary Figure 5

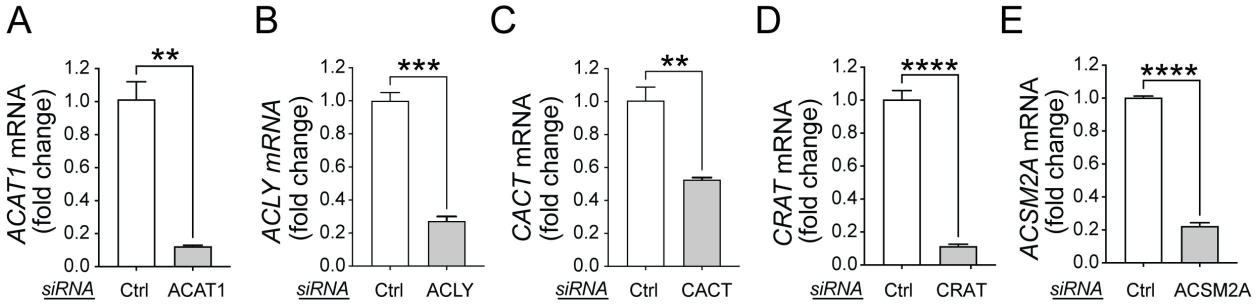


**Figure S5. Validation of various genes' siRNA knockdown by qRT-PCR.**

mRNA levels of ACAT1, ACLY, CACT, CRAT and ACSM2A were reduced by over 50% in human monocytes with corresponding siRNA nucleofection as compared to control siRNA nucleofection. Data were analyzed by unpaired two-tailed Student's *t*-test. All data were represented as mean ± SEM. **$p<0.01$; ***$p<0.001$; ****$p<0.0001$.
